# Role of the IRE1α-XBP1 axis in IgE-dependent activation of mast cells

**DOI:** 10.64898/2026.02.17.706498

**Authors:** Hiroto Kouda, Kazuki Nagata, Riu Saito, Chiharu Nishiyama

## Abstract

The IRE1α-XBP1 axis is the most conserved of three major unfolded protein response (UPR) branches that are triggered by the endoplasmic reticulum (ER) stress. Although the transcription factor XBP1 is involved in the development and function of several hematopoietic lineages, the role of XBP1 in the activation of mast cells (MCs) that play key role in allergic response remains largely unknown. Because we have identified salicylaldehyde (SA), which inhibits IRE1α nuclease activity that is essential for production of XBP1, as an inhibitor of MC activation in our previous screening, we investigated the effects of additional IRE1α inhibitors, 3-methyl-6-bromo-salichylaldehyde (MBSA) and KIRA6, targeting nuclease domain and kinase domain, respectively, on MC activation. MBSA and KIRA6 suppressed IgE-dependent degranulation and cytokine release of bone marrow-derived MCs (BMMCs), whereas these inhibitors did not suppress the Ca^2+^ ionophore- or compound48/80-induced degranulation. Treatment with inhibitors against two other branches of UPR, the PERK and the ATF6 pathways, did not affect IgE-induced activation of BMMCs. Intraperitoneal administration of MBSA or KIRA6 significantly suppressed IgE-induced passive anaphylaxis in mice. Furthermore, to evaluate the effect of XBP1, siRNA-mediated knockdown was performed. It was confirmed that *Xbp1* siRNA introduction reduced IgE-dependent degranulation of BMMCs in parallel with the knockdown level of *Xbp1* mRNA.

Taken together, the IRE1α-XBP1 axis plays a significant role in IgE-dependent and MC-mediated allergic response, which is considered to be therapeutic target of allergic diseases.

## Introduction

Mast cells (MCs) play important roles in IgE-mediated immune responses, including pathogenesis of allergic diseases and defense against parasite infection. In the case of allergic patients carrying hyper IgE, FcεRI expressed on MCs are occupied with antigen specific IgE, and cross-linking of multiple IgE on FcεRI with allergens rapidly activates MCs, resulting in degranulation, eicosanoid release, and cytokine production. In addition, several receptors, such as ST2, MRGPRX2, TLR4, and P2X7, which recognize IL-33, chemical compound, LPS, and ATP, respectively, are involved in IgE-independent activation of MCs. Current anti-allergic drugs, such as inhibitors of histamine receptor, leukotriene receptor, and anti-human IgE, target the events relating to MC activation, demonstrating that regulation of MC activation is effective for prevention of allergic diseases ^1^.

We have previously performed a screening to identify novel anti-allergic compounds using mouse bone marrow-derived MCs, and selected salicylaldehyde (SA) as the most effective inhibitor of MC activation^2^. The administration of SA suppressed IgE-dependent passive anaphylaxis in mice, although the molecular mechanisms by which SA inhibited IgE-dependent activation of MCs in vivo and in vitro remained unclear.

In the present study, we investigated the effects of the IRE1α-XBP1 axis that is the most conserved arm of the unfolded protein response (UPR) in MC activation, because this axis was considered the target of SA ^3–5^. The UPR occurred in response to the endoplasmic reticulum (ER) stress that is induced by accumulation of misfolded proteins within the ER lumen. Following events are induced under the UPR condition to maintain cellular homeostasis: i.e., degradation of unfolded proteins is promoted, further protein synthesis is inhibited, and repair mechanisms is facilitated. IRE1α is an enzyme possessing both kinase and endoribonuclease activities, and homodimerized-and phosphorylated-IRE1α creates the spliced mature *XBP1* mRNA by cleaving out 26 nucleotides, resulting in the translation of the transcription factor XBP1, which upregulates UPR-related genes. XBP1 also regulates several cellular events, such as protein secretion, lipid biosynthesis, glycosylation, and autophagy ^6^. XBP1 is involved in various immune system functions, including differentiation of plasma cells ^7^, development and survival of dendritic cells ^8^ and eosinophils ^9^, TLR-mediated cytokine production of macrophages ^10^, regulation of type I interferon production in chronically activated pDC ^11^, and anti-tumor activity of dendritic cells ^12^. On the other hand, the role of XBP1 and/or UPR in MC activation associated with cellular stresses remains largely unexplored. Although quite recently several reports regarding the role of XBP1 in MC function were published, they are still controversial ^13–15^ (Ref.^13^ has been corrected in Ref.^14^). Our study demonstrated the positive role of the IRE1α-XBP1 axis in IgE-induced activation of MCs.

## Materials and Methods

### Mice and Cells

C57BL/6J male and female mice (Japan SLC, Hamamatsu, Japan) housed under specific pathogen-free conditions were utilized for passive anaphylaxis model (6-8 weeks old) or sacrificed to generate bone marrow-derived MCs (BMMCs) (6-10 weeks old). BMMCs were prepared under recombinant mouse IL-3 (BioLegend, San Diego, CA, USA) supplemented conditions as previously described ^16^.

### Reagents

Ant-TNP mouse IgE (clone IgE-3, BD Bioscience, San Jose, CA, USA), TNP-BSA (LSL, Tokyo, Japan), 4-nitrophenyl-*N*-acetyl-β-D-glucosaminide (#N0866, Tokyo Chemical Industry, Tokyo, Japan), salicylaldehyde (SA; #S0004, Tokyo Chemical Industry), 3-methyl-6-bromo-salichylaldehyde (MBSA; #B1487, Tokyo Chemical Industry), KIRA6 (#S8658, Selleck Chemicals, Houston, TX, USA), GSK2606414 (#T2614, TargetMol, Boston, MA, USA), Ceapin-A7 (#T9110, TargetMol), A23187 (#11016, Funakoshi, Tokyo, Japan), compound 48/80 (#C2313, Sigma-Aldrich, USA), Tunicamycin (#ab120296, Abcam, UK), and DMSO (07-4860-5, Sigma Aldrich) were purchased from the indicated sources.

### Degranulation of MCs

Degranulation assay was performed as previously described method^17^ by sensitizing BMMCs (5.0 × 10^5^) with 0.2 μg/mL anti-TNP mouse IgE in 1 mL medium at 37 °C for 2 h, and then stimulated by 3 ng/mL TNP-BSA in 200 μL Tyrode’s buffer after washing with Tyrode’s buffer. After 30 min stimulation, culture supernatants were harvested for the β-hexosaminidase assay ^17^.

### Quantification of mRNAs

RevaTraAce qPCR RT Master Mix (#FSQ-201, TOYOBO, Osaka, Japan) were used for extraction of total RNA and for synthesis of cDNA, respectively. Quantitative real-time PCR was performed on a StepOne Real-Time PCR System (Applied Biosystems) with THUNDERBIRD SYBR qPCR Mix (#QPS201, TOYOBO) using the following primers.

*Xbp1*u-F; 5’-GACAGAGAGTCAAACTAACGTGG-3’

*Xbp1*u-R; 5’-GTCCAGCAGGCAAGAAGGT-3’

*Xbp1*s-F; 5’-AAGAACACGCTTGGGAATGG-3’

*Xbp1*s-R; 5’-CTGCACCTGCTGCGGAC-3’

*Actb*-F; 5’-AGATGACCCAGATCATGTTTGAGA-3’

*Actb*-R; 5’-CACAGCCTGGATGGCTACGTA-3’

### ELISA

The concentrations of mouse IL-6 and TNF-α were determined by ELISA MAX Standard/Deluxe Set-series ELISA kits (#431316 for IL-6; #430901 and #430916 for TNF-α).

### Flow cytometric analysis

BMMCs pretreated with 1 μg/mL Fc block (clone 93, BioLegend) were stained with 2.5 μg/mL DAPI (#11034-56, Nacalai Tescue, Kyoto, Japan), PE anti-mouse FcεRI (clone MAR1, BioLegend, 1:500), FITC anti-mouse c-kit (clone 2B, BioLegend, 1:200), and APC anti-mouse c-kit (clone ACK2, BioLegend, 1:200). Fluorescence was detected by a FACSLyric Analyzed (BD Pharmingen) or a MACS Quant Analyzer (Miltenyi Biotec, Tubingen, Germany), and the data were processed with FlowJo software (Tomy Digital Biology, Tokyo, Japan).

### Transfection of siRNA

Small interfering RNAs for mouse *Xbp1*; *Xbp1*-1 (Xbp1-MSS278861), *Xbp1*-2 (Xbp1-MSS278862), *Xbp1*-3 (Xbp1-MSS278863), and Stealth RNAi siRNA Negative Control MedGC (#12935-300) were purchased from Invitrogen (Carlsbad, CA, USA). BMMCs (2.0 ∼ 6.0 × 10^6^) were transfected with 10 μl of 20 μmol/L siRNA by using Neon transfection system (Invitrogen) as previously described ^16^.

### Passive anaphylaxis

Passive systemic anaphylaxis (PSA) and passive cutaneous anaphylaxis (PCA) were introduced to C57BL/6 mice by administration of anti-TNP mouse IgE and TNP-BSA as described in our previous study ^18^. MBSA (50 ∼ 100 mg/kg), KIRA6 (2.5 mg/kg), or vehicle (saline:PEG:Tween-80:DMSO=10:8:1:1 for MBSA, saline:DMSO:Tween-80=18:1:1 for KIRA6) was administered via an intraperitoneal (i.p.) injection once per day.

### Statistical analysis

A two-tailed Student’s t-test was used to compare two samples. To compare more than three samples, a one-way ANOVA followed by Dunnett’s multiple comparison test was performed. *p*-Values <0.05 were considered to be significant.

## Results

### Effect of SA-related compound MBSA and an XBP1 splicing inhibitor Kira6 on IgE-induced degranulation of BMMCs

We have identified SA as an inhibitor of MC activation through the screening of a chemical/natural compound library in our previous study ^2^. Because SA has been used for inhibition of *Xbp1* mRNA splicing by IRE1α in keratinocyte^3^ and melanoma^4^, we herein investigated the role of XBP1 and XBP1-related cellular events including UPR in MC activation. First, we evaluated the effect of SA-related compound MBSA on MC activation, which possesses higher inhibitory activity on XBP1 splicing than that of SA^5^. As shown in **Fig. 1A left**, treatment with MBSA for 48 h suppressed IgE-induced degranulation of BMMCs even in lower dose than the case of SA, and the suppressive effect was observed under the condition that MBSA was added to culture media 2h before or just when FcεRI-mediated stimulation was induced (**Fig. 1A right**). Then, we investigated the involvement of IRE1α-XBP1 pathway in IgE-induced activation of MCs by using another inhibitor, KIRA6, which inhibits XBP1 activation by targeting kinase domain of IRE1α ^19^, and found that KIRA6 significantly suppressed IgE-induced degranulation (**Fig. 1B**).

**Fig. 1.**
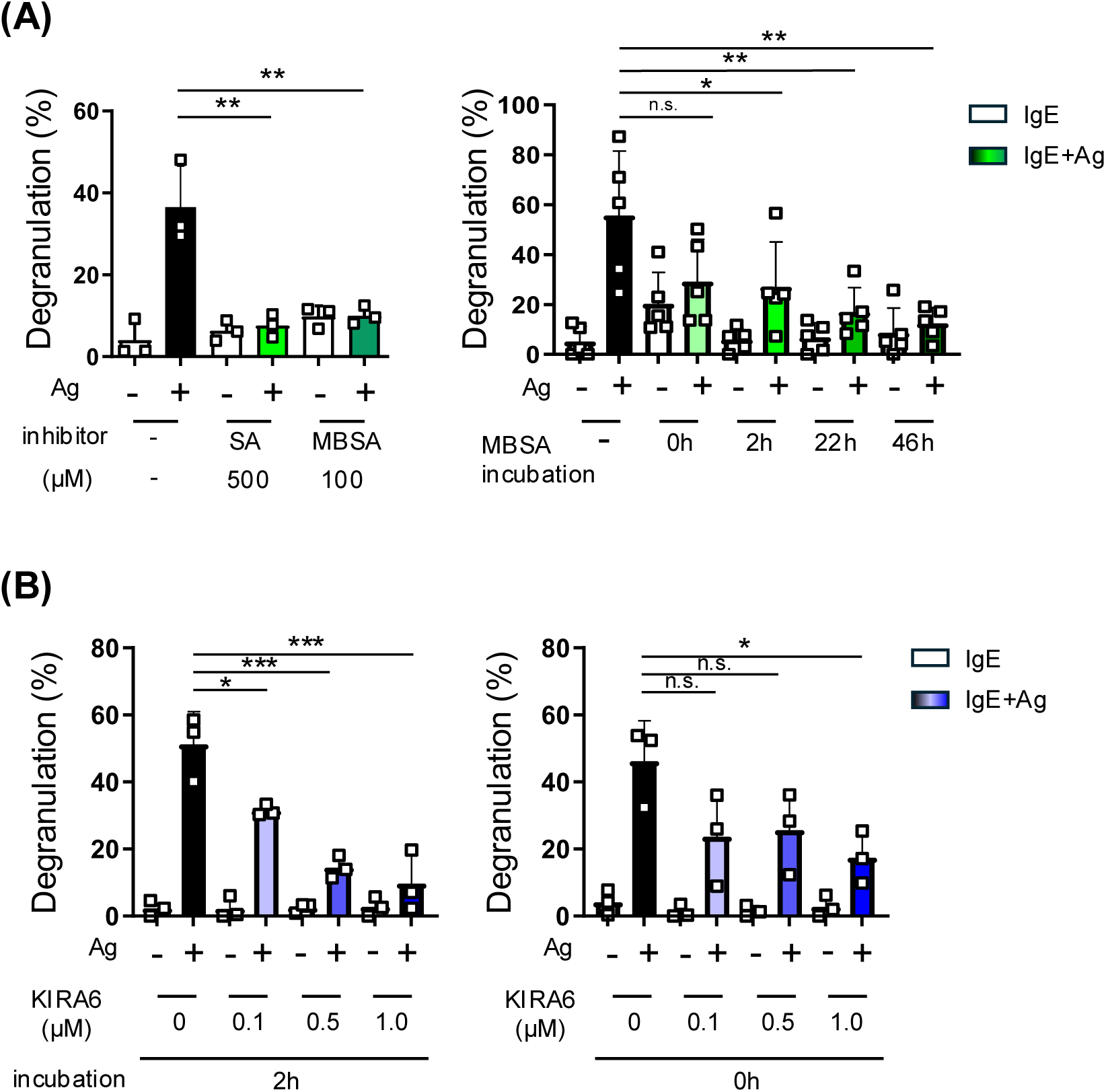
Effects of IRE1α inhibitors, SA, MBSA, and KIRA6, on IgE-induced degranulation of BMMCs. **A.** IgE-induced degranulation of BMMCs treated with SA or MBSA. Right; BMMCs were treated with 500 μM SA or 100 μM MBSA for 48 h, left; BMMCs were treated with 100 μM MBSA for indicated h. After treatment with SA or MBSA, BMMCs (5 × 10^5^ cells /mL) were incubated with 200 ng/mL of mouse IgE for 2h. After washing BMMCs with culture medium for two times, BMMCs suspended in Tyrode’s buffer were stimulated with 3 ng/mL of TNP-BSA as antigen (Ag) for 30 min. Beta-hexosaminidase activities in supernatants and in cells were measured as described in our previous study to determine degranulation degree. **B.** Inhibitory effect of KIRA6 on IgE-induced degranulation of BMMCs. Indicated concentrations of KIRA6 were added with IgE (left; 2 h before Ag stimulation) or just when BMMCs were stimulated with Ag (right). These experiments were repeated on different day with independently prepared BMMCs from different individuals. The data represent the mean ± SD of independent experiments. Dunnett’s multiple comparison test was used for statistical analysis between Ag stimulated BMMCs without inhibitor (Ctrl) and Ag stimulated BMMCs with an inhibitor. *; *p* < 0.05, **; *p* < 0.01.

These results suggested that inhibition of the IRE1α-XBP1 pathway suppressed IgE-induced degranulation of MCs.

### MBSA and KIRA6 suppressed IgE-induced cytokine production but did not affect IgE-independent degranulation of BMMCs

Upon IgE-dependent activation, MCs cause not only degranulation as the immediate reaction but also release of inflammatory cytokines involving the late phase reaction. To evaluate the effect of XBP1 inhibition on cytokine release, we determined the amount of IL-6 and TNF-α produced from IgE-stimulated BMMCs using ELISA. As the result, both treatment with MBSA (**Fig. 2A**) and KIRA6 (**Fig. 2B**) significantly suppressed the IgE-induced secretion of IL-6 and TNF-α from BMMCs.

**Fig. 2.**
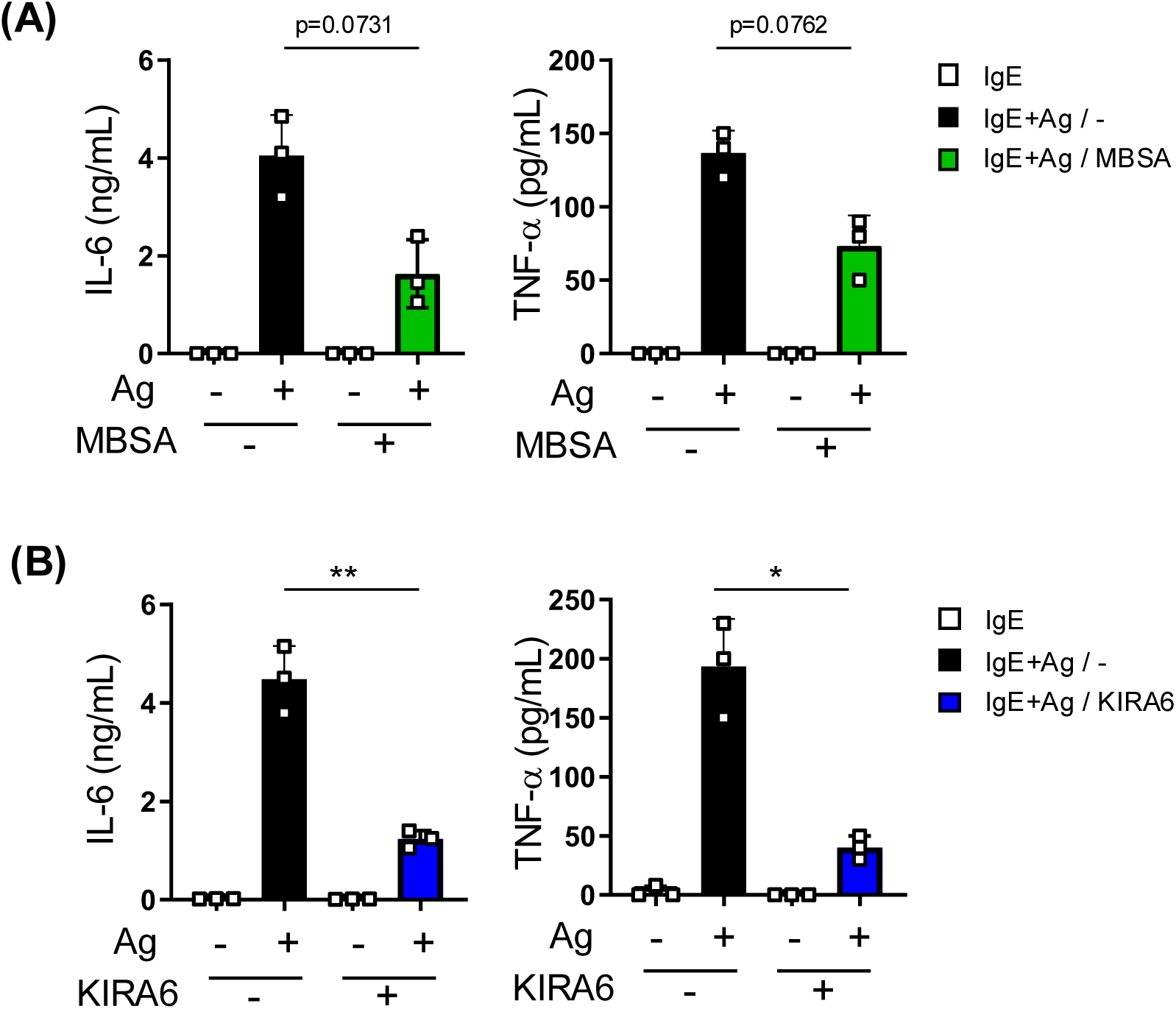
Effects of MBSA and KIRA6 on IgE-induced cytokine secretion from BMMCs. Concentrations of IL-6 (left) and TNF-α (right) in culture supernatants of IgE-stimulated BMMCs with or without treatment of MBSA (**A**) and KIRA6 (**B**). After 2 h incubation of BMMCs (5 × 10^5^ cells /mL) with 200 ng/mL of mouse IgE, BMMCs were washed with culture medium, and were then stimulated with 3 ng/mL of TNP-BSA as antigen (Ag). Culture supernatants were harvested 3 h after Ag-stimulation for ELISA. MBSA (final concentration 100 μM) was added at 48 h before Ag stimulation, and KIRA6 (1 μM) was added just before the addition of Ag. These experiments were repeated on different day with independently prepared BMMCs from different individuals. The data represent the mean ± SD of independent experiments. Two-tailed paired Student’s *t*-test was used for statistical analysis between Ag stimulated BMMCs without inhibitor (−) and Ag stimulated BMMCs with an inhibitor (+). *; *p* < 0.05, **; *p* < 0.01.

Furthermore, we investigated the effects of inhibitors on IgE-independent stimulation of BMMCs. Using a β-hexosaminidase assay, we found that compound 48/80-induced degranulation was not affected by pretreatment with MBSA, whereas Ca^2+^ ionophore-induced degranulation was significantly inhibited by MBSA (**Fig. 3A**). In contrast, KIRA6 did not suppress these IgE-independent degranulation of BMMCs (**Fig. 3B**).

**Fig. 3.**
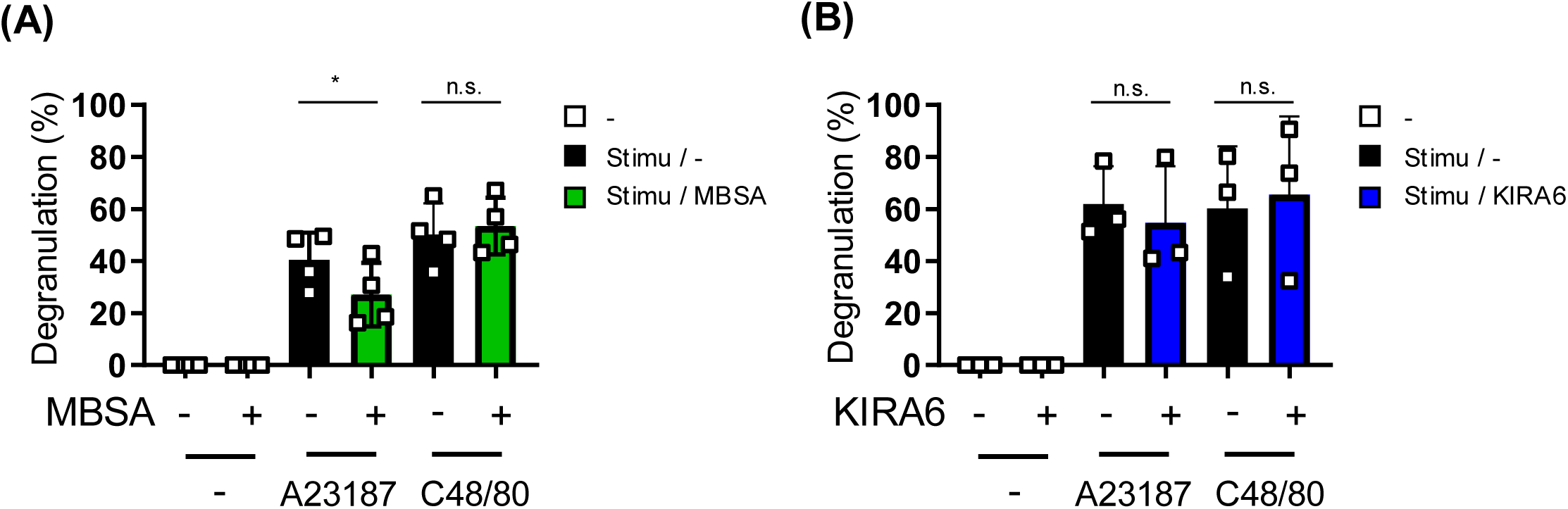
MBSA and KIRA6 suppresses Ca^2+^ ionophore- and compound 48/80-induced degranulation of BMMCs. Degranulation levels of BMMCs stimulated by A23187 (left) or compound 48/80 (C48/80; right) with or without treatment of MBSA (**A**) and KIRA6 (**B**). BMMCs incubated with 100 μM MBSA for 24 h were stimulated by 1 μM of A23187 (left) or 30 μg/mL of C48/80 (right) for 30 min (**A**). One μM KIRA6 was added just before the addition of A23187 or C48/80 (**B**). The harvested supernatants and collected cells were assessed to β-hexosaminidase assay (see Materials and Methods). These experiments were repeated on different day with independently prepared BMMCs from different individuals. The data represent the mean ± SD of independent experiments. Two-tailed paired Student’s *t*-test was used for statistical analysis between without inhibitor (−) and with an inhibitor (+). *; *p* < 0.05, n.s.; not significant.

### Analysis of involvement of other two UPR pathways on MC activation by using specific inhibitors

In addition to the IRE1α-XBP1, the PERK-ATF4 and the ATF6 pathways are involved in the UPR. Then, we evaluated the effects of GSK2606414 and CeapinA-7, which inhibit the activation of PERK ^20^ and the trafficking of ATF6 ^21^, respectively, on MC activation. We pretreated BMMCs with 0.1 – 10 μM of GSK2606414 or CeapinA-7 for 24 h based on previously reported experimental conditions. As results, both GSK2606414 (**Fig. 4A**) and CeapinA-7 (**Fig. 4B**) did not exhibit apparent effects on IgE-, A23187-, and Compound 48/40-induced degranulation of BMMCs.

**Fig. 4.**
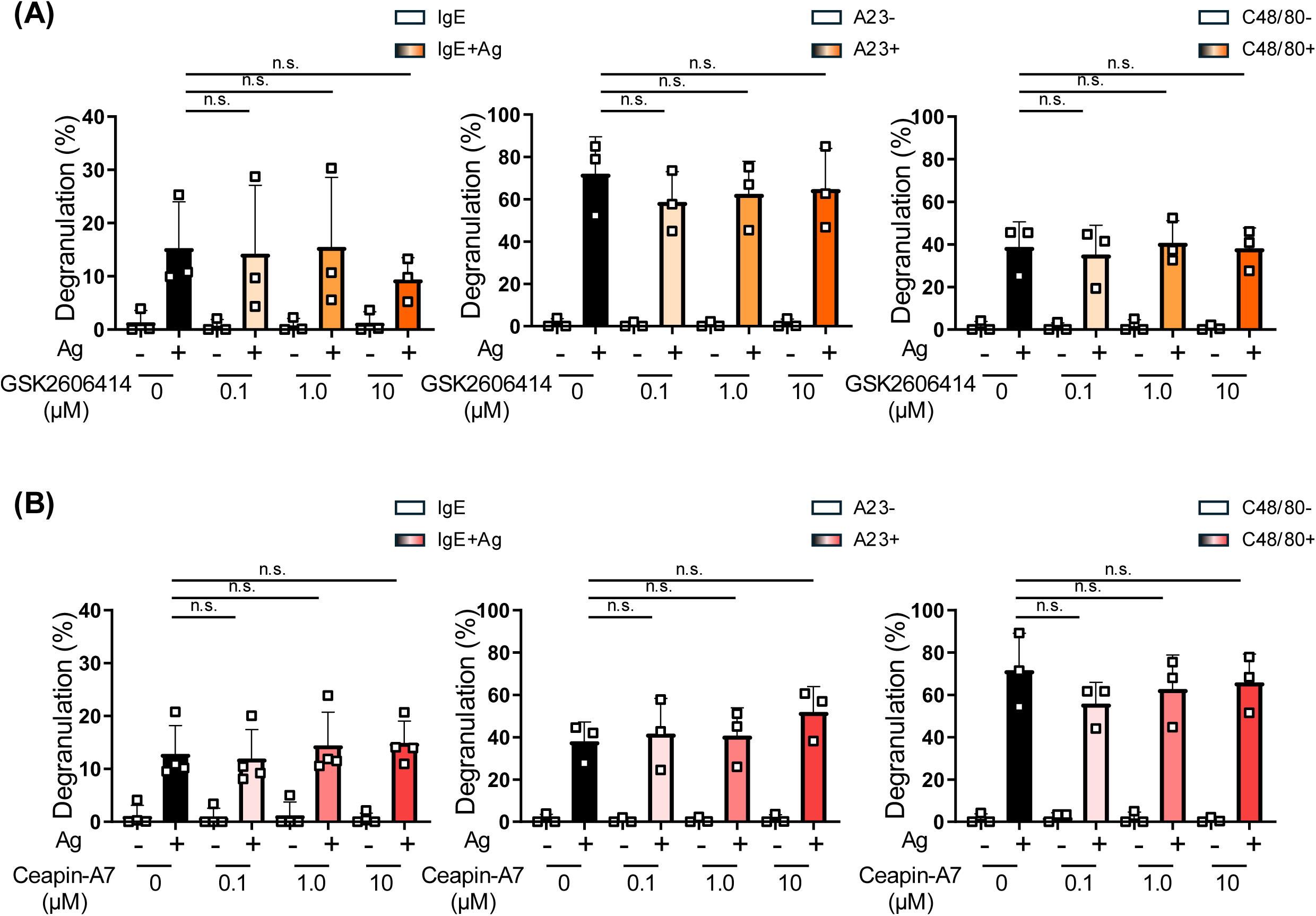
Inhibitors targeting two other pathways did not affect activation of BMMCs. BMMCs were incubated with indicated concentrations of GSK2606414, a PERK inhibitor (**A**), or CeapinA-7, an ATF6 inhibitor (**B**), for 22 h before the addition of IgE and for additional 2 h during incubation with Ag (left), or for 24 h before the addition of A23187 (center), and compound 48/80 (right). These experiments were repeated on different day with independently prepared BMMCs from different individuals. The data represent the mean ± SD of independent experiments. Two-tailed Dunnett’s multiple comparison test was used for statistical analysis between without inhibitor (Ctrl) and with an inhibitor. n.s.; not significant.

These results suggest the IRE1α-XBP1-dependent and the PERK-ATF4/ATF6-independent manner of IgE-induced activation of MCs as observed in other events relating to drastic protein synthesis and immuno-responses ^7,22,23^.

### Effects of Tunicamycin on MC activation

Tunicamycin, which inhibits *N*-linked glycosylation of protein, causes accumulation of unfolded proteins in ER and subsequent ER stress. In a recent study reporting the association of ER stress with MC degranulation, tunicamycin i.p. administration raised the amount of serum Mcpt1 in mice ^13,14^, which is a hallmark of MC activation. Then, we treated BMMCs with tunicamycin to investigate whether induction of ER stress upregulates the IgE-induced activation of MCs. As shown in **Fig. 5A**, tunicamycin pretreatment suppressed IgE-induced degranulation of BMMCs in a dose-dependent manner. The treatment of BMMCs with 10 μg/mL tunicamycin, which reduced the degranulation significantly, completely inhibited IgE-dependent secretion of IL-6 and TNF-α (**Fig. 5B**). We also revealed that surface expression of FcεRI on BMMCs was markedly reduced by the treatment with tunicamycin (**Fig. 5C**).

**Fig. 5.**
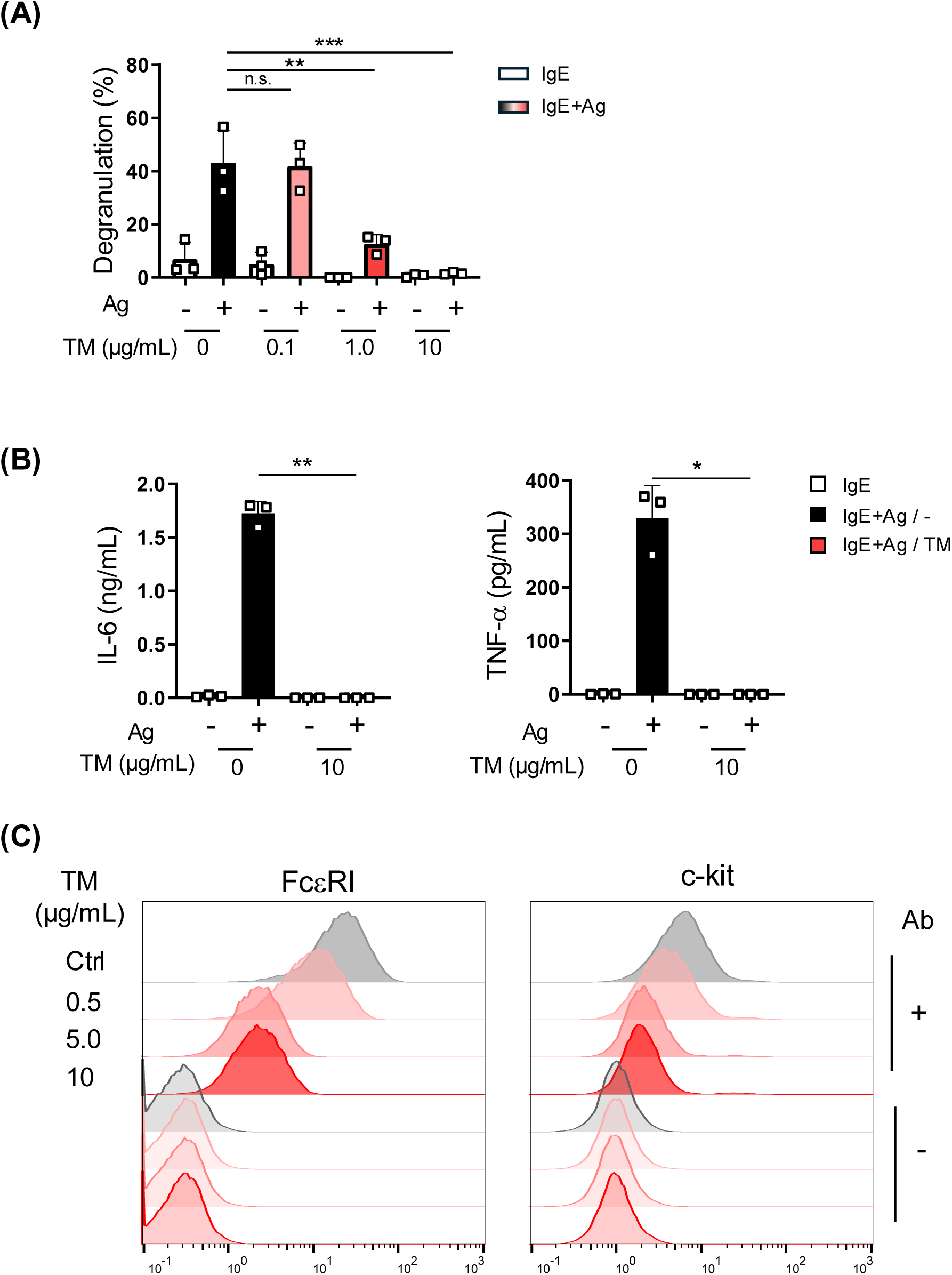
Effects of tunicamycin on MC activation. **A.** IgE-induced degranulation of BMMCs treated with tunicamycin. **B.** IL-6 (left) and TNF-α (right) production from tunicamycin-treated BMMCs. **C.** Cell surface expression of FcεRI (right) and c-kit (left) on tunicamycin-treated BMMCs. BMMCs (5 × 10^5^ cells /mL) were incubated in the presence or absence of indicated concentration of tunicamycin (TM) for 22 h, and then were subjected to IgE-induced degranulation assay (**A**), IgE-induced cytokine production (**B**), and to flow cytometric analysis (**C**). These experiments were repeated on different day with independently prepared BMMCs from different individuals. The data represent the mean ± SD of independent experiments. Dunnett’s multiple comparison test was used in (**A**) and two-tailed paired Student’s *t*-test was used for statistical analysis between without TM (Ctrl) and with TM (**B**).

Since *N*-glycosylation is required for multiple processes in MC function, including the transport of FcεRI from the ER to the cell surface ^24^, these results suggested that tunicamycin does not specifically induce the UPR as predicted in a previous study, but inhibited MC activation by instead exerts a broad effect on MCs ^13^.

### MBSA and KIRA6 suppressed IgE-dependent anaphylaxis in mice

Above-mentioned results of in vitro experiments using BMMCs demonstrated that the inhibitors for the IRE1α-XBP1 pathway suppressed IgE-induced activation of MCs. To evaluate the effects of these inhibitors on IgE-dependent allergic response in vivo, we utilized passive systemic anaphylaxis models. As shown in **Fig. 6A**, i.p. administration of MBSA significantly suppressed swelling of the ear caused by i.v. injection of TNP-BSA into IgE-preinjected mice in PCA model. We also confirmed that body temperature drop induced upon antigen injection in PSA model was reduced in mice administered MBSA or KIRA6 (**Fig. 6B**).

**Fig. 6.**
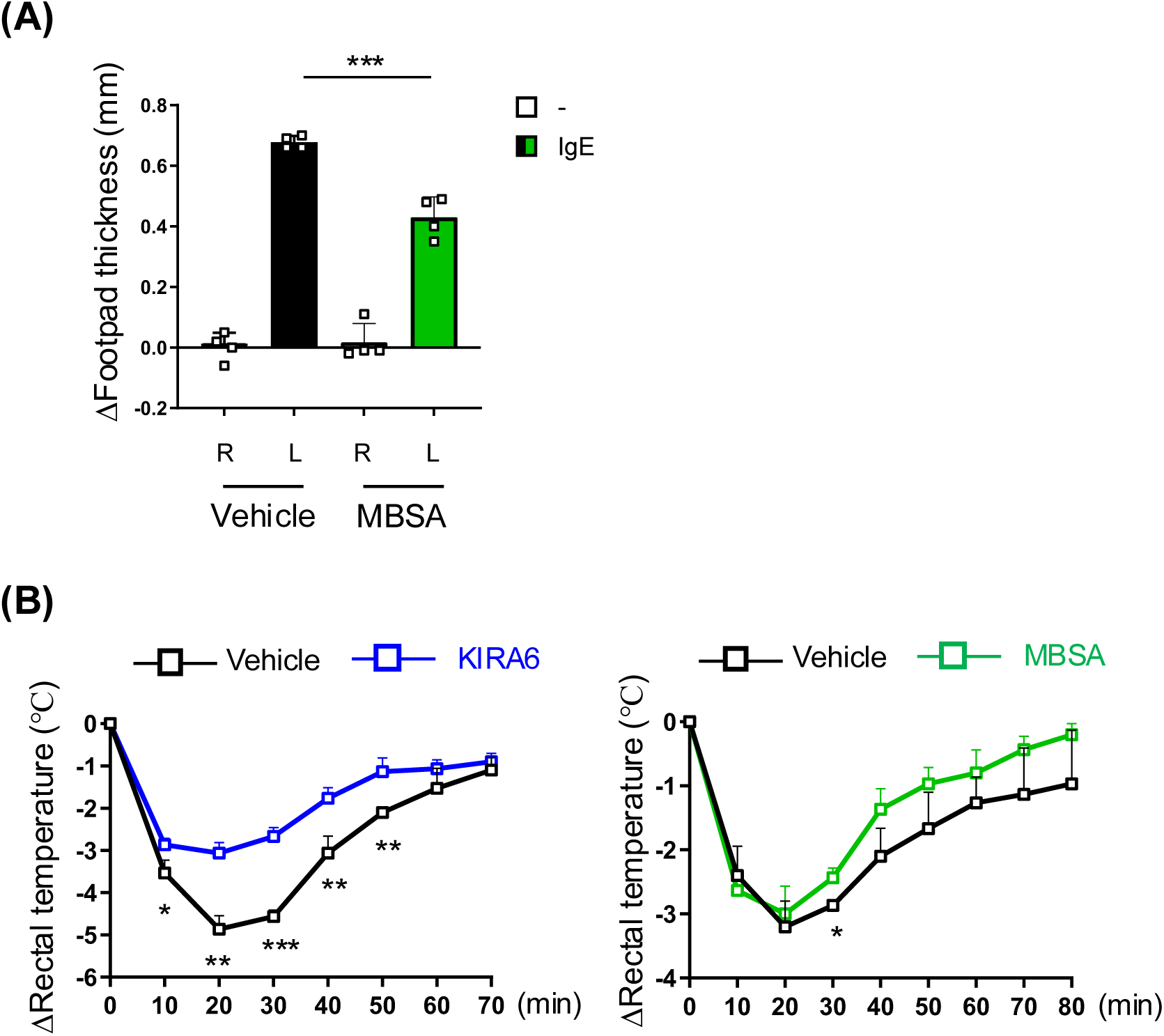
Passive anaphylaxis levels of mice administered MBSA or KIRA6. **A.** Footpad swelling of passive cutaneous anaphylaxis model mice. Mice were i.p. injected with MBSA (50 mg/kg/day) or 200 μl vehicle (Ctrl; see Materials and Methods) for 5 days (n = 4 for each group). IgE solution and its vehicle were injected into the left (L) and right (R) footpad, respectively, on day4, and Ag was i.v. injected on day5. Footpad thickness 30 min after Ag injection was measured as described in our previous study ^2,18^. Unpaired Student’s *t*-test was used for statistical analysis between Ctrl and MBSA. ***; *p* < 0.005. **B.** Body temperature of passive systemic anaphylaxis model mice. Mice were i.p. injected with 100 mg/kg/day MBSA or its vehicle (right), and 2.5 mg/kg/day KIRA6 (right) or its vehicle (left) for 3 days (n = 3 for each group). I.v. injections of IgE on day2 and of Ag on day3 were performed as described previously ^2,18^. The data represent the mean ± SD of independent experiments. The unpaired Student’s *t*-test was used for statistical analysis between vehicle and inhibitor. *; *p* < 0.05.

These results indicated the biological significance of the IRE1α-XBP1 pathway in the IgE-dependent anaphylaxis.

### XBP1 is involved in IgE-induced responses of MCs

Quite recently, 2 reports regarding the effects of KIRA6 on MC activation have been published ^15,25^. These 2 studies claimed that KIRA6 suppressed MC activation via inhibiting the kinases at downstream of FcεRI, such as Lyn and/or Fyn, rather than IRE1α. These conclusions prompted us to analyze the involvement of XBP1 itself in IgE-induced activation of MCs. To clarify the role of XBP1 in MC activation, we performed knockdown experiment using siRNA transfection. Among three clones of *Xbp1* siRNAs, *Xbp1*-2 siRNA most effectively reduced mRNA levels of spliced *Xbp1* (*Xbp1s*), whereas the knockdown efficiency of *Xbp1*-1 and -3 siRNAs were moderate (**Fig. 7A**). Beta-hexosaminidase assay of IgE-dependently activated BMMCs revealed that degranulation degree of *Xbp1* siRNA-transfected BMMCs decreased in parallel with knocked down levels of *Xbp1* mRNA (**Fig. 7B**). We confirmed that levels of *Xbp1* mRNA reduction (**Fig. 7C**) and degranulation suppression (**Fig. 7D**) by *Xbp1*-2 siRNA were statistically significant.

**Fig. 7.**
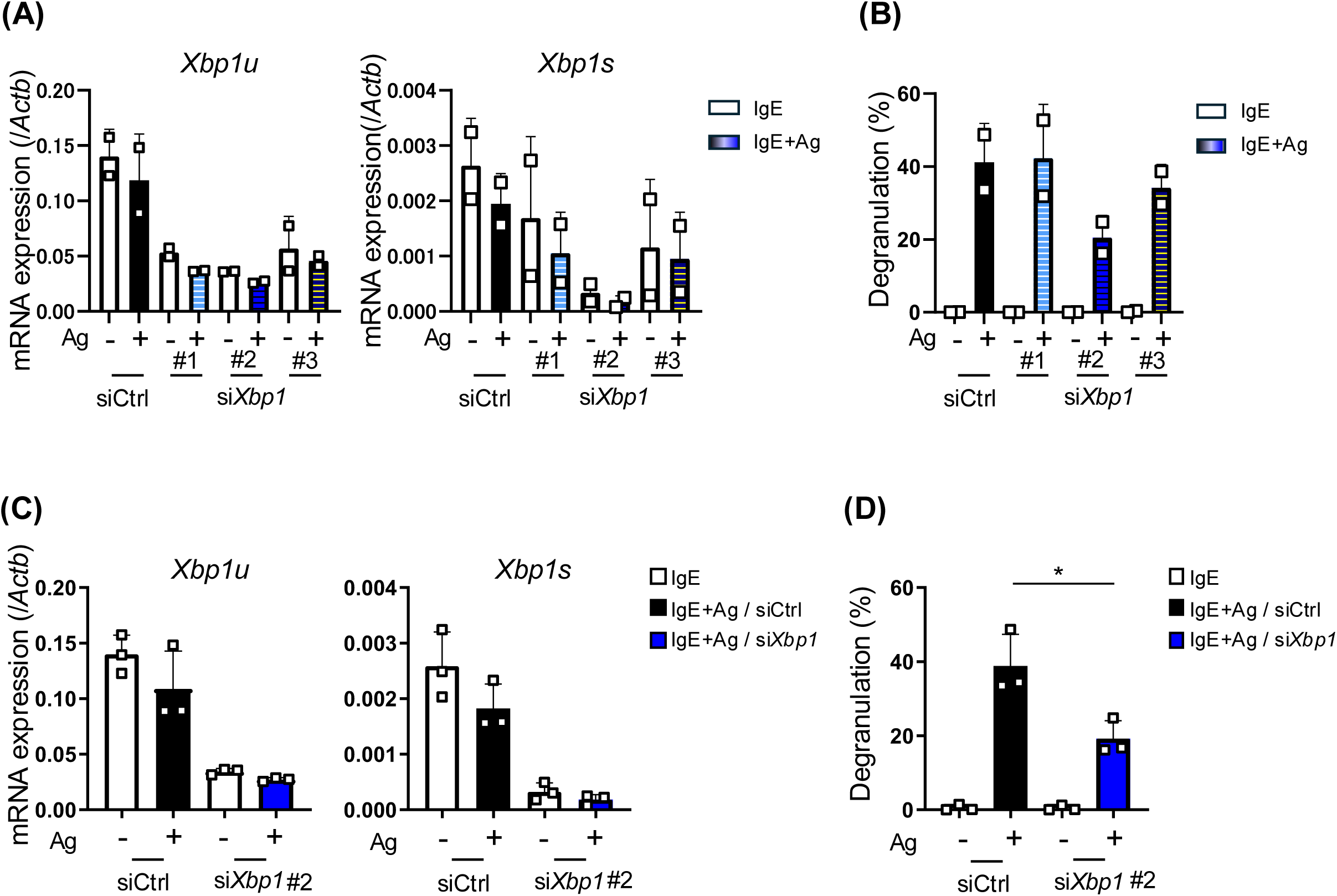
Effects of *Xbp1* siRNA introduction on IgE-induced activation of BMMCs. **A.** and **C.** mRNA levels of un-spliced *Xbp1* (*Xbp1u*) and spliced *Xbp1* (*Xbp1s*) in siRNA transfected BMMCs, which were harvested 3 h after Ag stimulation. **B.** and **D.** Degranulation degree of *Xbp1* siRNA-transfected BMMCs. After 72 h culture from electroporation, BMMCs were divided into two groups: one for mRNA quantification and the other for degranulation assay. Three different sequences of *Xbp1* siRNAs were used: *Xbp1*-1 (Xbp1-MSS278861), *Xbp1*-2 (Xbp1-MSS278862), *Xbp1*-3 (Xbp1-MSS278863). These experiments were repeated on different day with independently prepared BMMCs from different individuals. The data represent the mean ± SD of independent experiments. The two-tailed paired Student’s *t*-test was used for statistical analysis between control siRNA-transfected BMMCs (*siCtrl*) and *Xbp1* siRNA-transfected BMMCs (*siXbp1*). *; *p* < 0.05.

Taken these results together, we demonstrated that XBP1 is required for IgE-dependent activation of MCs.

## Discussion

We have previously identified SA as the most effective compound that inhibited the IgE-dependent activation of MCs in a screening study aiming the development of novel anti-allergic drug^2^. Since SA is reported to inhibit the splicing of *Xbp1* mRNA by IRE1α ^3,4^, which is required for the translation of functional XBP1 protein, we investigated the involvement of the IRE1α-XBP1 axis in IgE-dependent activation of MCs in the present study. We revealed that two other IRE1α inhibitors, SA-related compound MBSA ^5^ and KIRA6 ^19^, against the endoribonuclease activity and kinase activity, respectively, suppressed IgE-dependent activation of MCs in vivo and in vitro, whereas the inhibitors targeting PERK and ATF6, which are also involved in the UPR as well as IRE1α-XBP1, did not affect the activation of MCs. Furthermore, knockdown of *Xbp1* mRNA reduced IgE-dependent degranulation of MCs. From these results, we concluded that XBP1 plays a positive role in IgE-dependent activation of MCs, which would be a therapeutic target of allergic diseases.

XBP1 transactivates the genes relating to secretion of proteins and lipid mediators, including insulin processing protease in the β cells ^23^, antibody production in plasma cells ^7^, and prostaglandin-synthesis related enzymes in dendritic cells ^26^. Therefore, the genes expressed in activated MCs, which secrete various mediators such as proteases, eicosanoids, and cytokines, could be possible targets of XBP1. We are going to conduct further detailed analysis revealing the target genes of XBP1 in MCs.

During preparation of this manuscript, two studies reporting the anti-allergic effects of KIRA6 have been published ^15,25^. *Wunderle* et al. demonstrated that KIRA6 suppressed Ag-stimulated signaling of MCs by binding to kinases, LYN, FYN, and KIT ^25^. The other report claimed that KIRA6 inhibited the kinase activity of Lyn in an IRE1α-independent manner because knock out of the *Ern1* gene (encoding IRE1α) by using genome editing did not affect Ag-induced activation of RBL-2H3, a rat basophilic cell line ^15^. These studies prompted us to perform knock down of *Xbp1* in BMMCs to reveal the role of XBP1 in IgE-dependent activation of BMMCs, because the experiment revealing direct effect of XBP1 is required and primary cells rather than leukemia cells should be used. Our result showing that degranulation degree of BMMCs decreased in parallel with *Xbp1* mRNA level indicated the positive role of XBP1 in the activation of MCs. Several previous studies regarding conditional KO of *Xbp1* demonstrated the requirement of XBP1 in the development and specific gene expression of hematopoietic cells ^8,26,27^. XBP1 maintains the function of immune cells, which is not restricted in the UPR.

In a recent study, the effects of tunicamycin on MC function were investigated ^13^. This report showed that i.p. injection of tunicamycin for 3 days increased serum concentration of Mcpt1 in mice without conducting any allergic model, and that exposure of BMMCs to tunicamycin for 3 h increased phosphorylation of kinases, Lyn, Syk, and PLCγ1. From these results, it was concluded that ER stress induction by tunicamycin activated MCs. Since the effect of tunicamycin treatment on IgE-induced activation of MCs has not been examined in this study, it is unclear whether enhanced phosphorylation of kinases leads higher activation of MCs and which kind of stimulation induced Mcpt1 secretion from MCs in tunicamycin-treated mice. However, under our experimental conditions, tunicamycin treatment markedly suppressed IgE-dependent activation of BMMCs (Fig. 5). This result contrasted with our prediction, based on the previous study, that tunicamycin would enhance IgE-dependent activation of MCs by inducing ER stress. Drastic reduction in cell surface FcεRI levels in tunicamycin-treated BMMCs is considered the primary cause of the responsiveness defect to FcεRI-mediated activation. However, we also recognize that the block of protein glycosylation by tunicamycin broadly affected MC activation-related events, because A23187-induced activation was also suppressed in tunicamycin-treated BMMCs (data not shown).

In the present study, we treated BMMCs with IRE1α inhibitors for 0∼48 h incubation time, and found that MBSA exhibited higher inhibitory effects with longer incubation time, whereas KIRA6 added just before stimulation was most effective for suppression of MC activation. This difference may partly reflect the off-target effect of KIRA6 against kinases, Lyn and/or Fyn ^15,25^. In these studies, KIRA6 added 30 min^28^ or 1 h^25^ before stimulation inhibited IgE-dependent activation of MCs. The stronger inhibitory effect of KIRA6 than that of MBSA observed in PSA under our experimental conditions may be due to the additional target sides of KIRA6. However, we do not exclude the role of KIRA6 against IRE1α, because *Xbp1s* mRNA levels in Ag-stimulated BMMCs tended to be reduced in the presence of 1 μM KIRA6 ^25^. Our preliminary experiment showed that mRNA levels of *Xbp1s* in BMMCs decreased 2 to 48 h after MBSA treatment (data not shown); however, further experiments are needed to clarify how *Xbp1s* mRNA levels change over time following KIRA6 treatment under our experimental conditions. To evaluate the IRE1a-XBP1-dependency of the inhibitory effects of MBSA and KIRA6, experiments could be conducted using MCs lacking XBP1. This point could potentially be resolved by performing a degranulation assay of BMMCs generated from *Xbp1*-deficient mice to determine whether MBSA and KIRA6 exert additional inhibitory effects on *Xbp1-*knockout BMMCs. Furthermore, the use of *Xbp1-*knockout MCs may demonstrate that XBP1 is involved in the inhibitory effect of MBSA on A23187-induced MC activation.

The involvement of the IRE1α-XBP1 axis in the IgE-dependent activation of MCs has been controversial: an IRE1α deficiency did not affect the IgE-induced activation of RBL-2H3 cells ^15^, whereas IgE-dependent Syk phosphorylation and calcium mobilization were completely inhibited in *Xbp1* KO mouse MCs ^13^. The results obtained using IRE1α inhibitors and *Xbp1*-targeting siRNA suggest that this pathway plays a positive role in IgE-dependent activation; however, further studies are needed to resolve this issue. A previous study using *Ern1*-knockout BM dendritic cells have suggested that IRE1a-XBP1 signaling plays an essential role in prostaglandin biosynthesis ^26^. Furthermore, we have demonstrated that prostanoids, particularly PGE_2_, suppressed IgE-dependent activation of MCs ^18,29^. Recently, we observed increased levels of *Ptgs* mRNAs in MBSA-treated or *Xbp1* siRNA-transfected BMMCs (data not shown). It may be interesting to evaluate the involvement of XBP1 in PG-related genes in MCs. We are currently attempting to identify the target gene(s) of XBP1 in MCs, which we expect to provide insights into the role of XBP1 in MC activation.

Finally, it is important to note that because XBP1 is involved in multiple processes across various organs and cells, it is necessary to consider the potential side effects that may arise from systemic administration of the inhibitors of XBP1. Identifying the target gene(s) of XBP1 in MCs could lead to development of specific treatments for allergic diseases.

## Acknowledgments

We thank members of the Laboratory of Molecular Biology and Immunology, Department of Biological Science and Technology, Tokyo University of Science for their constructive discussions and technical support.

This work was supported by a Grants-in-Aid for Scientific Research (B) 23K26860 (CN), and 23H02167 (CN); a Grant-in-Aid for Early-Career Scientists 24K17872 (KN); a Tokyo University of Science Grant for President’s Research Promotion (CN); the Tojuro Iijima Foundation for Food Science and Technology (CN); a Research Grant from the Mishima Kaiun Memorial Foundation (CN); and a Research Grant from the Takeda Science Foundation (CN).

We greatly appreciate the consideration from Ms. Yayoi Yasuda, Dr. Masako Yasuda, and Dr. Kimihiko Yasuda.

## Author contributions

Conceptualization, C.N.; investigation - the majority of experiments and data analysis were performed by H.K. with assistance from K.N.; writing - original draft preparation, review, and editing, C.N.; supervision, C.N.; project administration, K.N., and C.N.; funding acquisition, K.N., and C.N. All authors have read and agreed to the published version of the manuscript.

## Supplemental information

None.

